# Mathematical modeling links benefits of short and long antibiotic treatment to details of infection

**DOI:** 10.1101/555334

**Authors:** Francisco F. S. Paupério, Vitaly V. Ganusov, Erida Gjini

**Affiliations:** Instituto Gulbenkian de Ciência, Rua da Quinta Grande, 6, 2780-156 Oeiras, Portugal; Department of Microbiology, University of Tennessee, Knoxville, TN 37996, USA; Departamento de Informática, Faculdade de Ciências, Universidade de Lisboa, Campo Grande, 1749-016, Lisbon, Portugal

**Keywords:** mathematical model, antibiotic therapy, antibiotic resistance, immunity

## Abstract

Antibiotics are the major tool for treating bacterial infections. With rising antibiotic resistance in microbes, strategies that limit further evolution and spread of drug resistance are urgently needed, in individuals and populations. While classical recommendations favor longer and aggressive treatments, more recent studies and clinical trials advocate for moderate regimens. In this debate, two axes of aggressive treatment have typically been conflated: treatment intensity and treatment duration, the latter being rarely addressed by mathematical models. Here, by using a simple mathematical model of a generic bacterial infection, controlled by host’s immune response, we investigate the role of treatment timing and antibiotic efficacy in determining optimal duration of treatment. We show that even in such simple mathematical model, it is impossible to select for universally optimal treatment duration. In particular, short (3 day) or long (7 day) treatments may be both beneficial depending on treatment onset, on the criterion used, and on the antibiotic efficacy. This results from the dynamic trade-off between immunity and resistance in acute, self-limiting infections, and uncertainty relating symptoms to the start of infection. We find that treatment timing can shift the trend between resistance selection and length of antibiotic exposure in individual hosts. We propose that major advances in predicting impact of antibiotics on bacterial infections must come from deeper experimental understanding of bacterial infection dynamics in humans. To guide rational therapy, mathematical models need to be constrained by data, including details of pathology and symptom thresholds in patients, and of host immune control of infection.

## Introduction

The treatment of bacterial infections has for many decades relied on the use of antibiotics. Although antibiotics have saved many lives and enabled uncountable medical practices, their widespread use in human and animal populations has led to the rise of antibiotic resistance, posing now a threat to human health and modern medicine [1]. Of particular concern is the rise of multidrug-resistant bacteria, favored by use of wide-spectrum antibiotics especially in clinical settings [2, 3]. To confront these challenges, much research has been devoted to understand the molecular, genetic, and non-genetic mechanisms leading to drug resistance in bacteria [4, 5, 6], their population dynamics, and interplay with treatment strategies [7, 8, 9]. While alternative approaches such as anti-virulence [10] or host-directed therapies [11] are also being considered, with their own potential limitations [12], reducing antibiotic use remains an essential step in addressing the antibiotic resistance crisis. In this context, it is important to understand the rational principles by which antibiotics succeed and fail in clearing infections, and whether and when aggressive or moderate treatments are superior.

The conventional wisdom of treating infections with high antibiotic doses (aggressive treatment) [13] to avoid resistance emergence has recently been challenged [14, 15], on the basis of evolutionary arguments showing a bigger risk of resistance selection in target pathogens with more aggressive treatments (see [9] for a review). Many studies including clinical trials have by now shown that for some infections shorter treatment is not inferior to the longer ones and that longer treatment may in fact result in failure if resistant bacteria are already present when treatment starts [16, 17, 18, 19, 20]. This issue is now recognized in clinical practice and checklists of improving antibiotic prescribing have been suggested [21].

On one hand, clinical studies have been concerned mainly with optimal duration of therapy, on the other, the multiple mathematical studies addressing the question of optimal antibiotic treatment of bacterial infections [22, 23, 24, 25], have focused mainly on the dosing dimension, with a few studies exploring duration [26] and timing of treatment [27]. While these studies have highlighted the various complexities in optimal treatment, typically two axes of aggression have been conflated: treatment length and treatment intensity, and a single criterion for defining optimality, e.g. resistance emergence or selection, has been considered.

Here, we extend these previous studies by formulating one of the simplest mathematical models of bacterial infection that is controlled by the immune response and investigate the role of treatment timing, intensity and duration, across different metrics of success. We show that duration of antibiotic treatment, antibiotic efficacy (defined as the antibiotic kill rate), and treatment timing interact nonlinearly to determine final outcome, and that optimal regimes vary widely with target criterion for optimization even at the single host level. Our results suggest that it is unlikely that one optimal treatment duration even exists, and that specific details of particular infections and antibiotic efficacy are important in determining the needed duration of treatment.

## Results

### The modeling framework

To investigate the impact of antibiotic treatment duration on treatment success or failure we utilized a mathematical model of a generic acute bacterial infection which includes the dynamics of three populations: drug-sensitive (*B*_*s*_) and drug-resistant (*B*_*r*_) bacteria and bacteria-specific immune response (*E*, see Eqns. (1)–(3) in Materials and Methods). Bacterial growth is described by a logistic equation with growth rate *r* and carrying capacity *K*, and the presence of bacteria induces immune response which expands exponentially and eventually clears the infection (Figure 1A). In our analysis, infections always start with drug-sensitive strain but at some level, a drug-resistant variant may appear due to mutations (at rate *m*); drug-resistance bears a fitness cost (*γ*). The proposed model is similar to previously published models [28, 23, 25]. In the model we assume that treatment starts when the total bacterial density *B* = *B*_*s*_ + *B*_*r*_ reaches a critical level Ω (symptom threshold). Treatment administration lasts for *τ* days with a given antibiotic efficacy (kill rate) *A*_*m*_ which represents the average net rate of antibiotic-induced bacterial killing at the infection site per unit of time.

**Figure 1:**
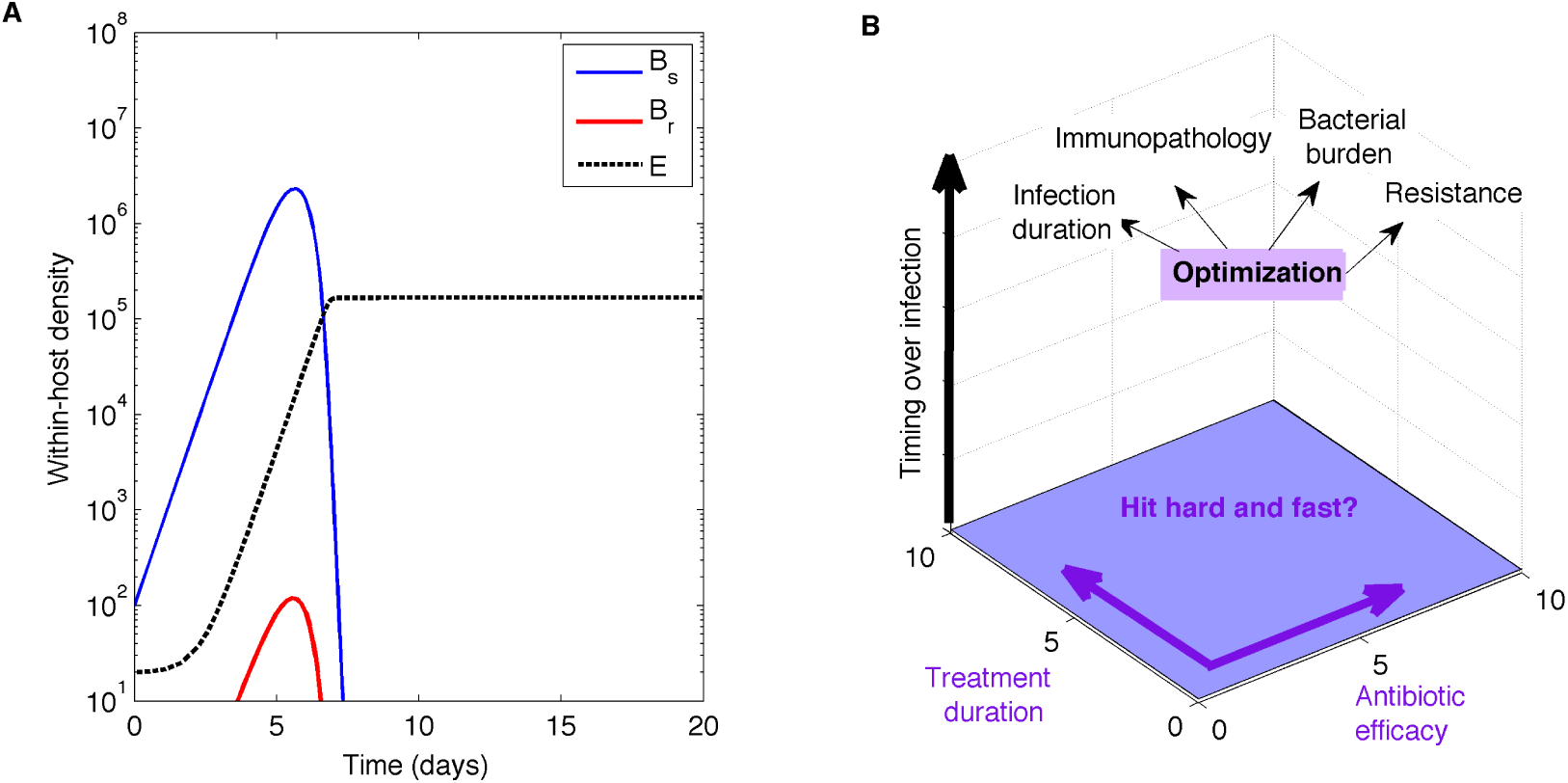
Typical dynamics of the bacterial infection in the mathematical model and treatment optimization challenge. **A**. Population dynamics of a generic bacterial infection with sensitive bacteria *B*_*s*_, emergence and potential selection of a drug-resistant sub-population *B*_*r*_ and host immune response *E* (see Eqns. (1)–(3)). Default parameters are given in Table 1. **B**. Treatment strategies evaluated in a 3-dimensional space (antibiotic efficacy, treatment duration and timing), along several target criteria of clinical and epidemiological importance.

Optimizing the duration of antibiotic treatment is likely to involve several alternative goals (e.g., reducing bacterial load, minimizing antibiotic resistance, speeding infection clearance [14, Figure 1B]), but from the perspective of the patient and treating physician, successful treatment generally means rapid reduction in symptoms and no disease relapse after treatment completion. To investigate whether alternative treatment goals conflict with the optimal duration of treatment, we track several instantaneous and cumulative measures such as the density of bacteria *B*(*t*) or the level of immunity *E*(*t*) at the end of treatment, the duration of infection, cumulative bacterial load (area under the curve, AUC_*B*_) and total resistance load (AUC_*R*_).

**Table 1:** Parameters of the mathematical model on bacterial infection. The mathematical model is given in Eqns. (1)–(3). We show the default set of parameters used in simulations as well as the range of parameters varied in some of our analyses. Most of these parameters have been chosen to give a reasonable dynamics for the bacteria and immune response (e.g., rapid bacterial and immune response growth). Yet, it should be noted that few if any of these parameters have been accurately measured for bacterial infections of humans..

| Symbol | Parameter | Default | Range |
| --- | --- | --- | --- |
| $r$ | Growth rate of bacteria | 2/day | |
| $K$ | Carrying capacity of bacteria | $10^7$ cell/ml | |
| $\gamma$ | Fitness cost of resistant bacteria | 0.1 | |
| $B_0$ | Initial inoculum | 100 cell/ml | |
| $\sigma$ | Maximal growth rate of the immune response | 2/day | |
| $k$ | Half-saturation constant for antigen-dependent immunity | $10^4$ cell/ml | $10^3 - 10^4$ |
| $d$ | Elimination rate of bacteria by host immunity | $1 \times 10^{-4}$/day | |
| $E_0$ | Initial immunity | 20 | |
| $m$ | Mutation rate conferring drug resistance per sensitive cell | $10^{-5}$ | |
| $B_{em}/B_{ext}$ | Resistance emergence/pathogen extinction threshold | 10 | |
| $P_{em}$ | Probability threshold for high-level resistance mutant | 0.5 | |
| $\Omega$ | Bacterial density at which treatment starts | $10^4$ cell/ml | $10^3 - 10^5$ |
| $A_m$ | Antibiotic kill rate of drug-sensitive bacteria (efficacy) | 1/day | 0.1 – 6/day |
| $\tau$ | Duration of treatment | 3 | 1 – 10 days |

### Both short and long treatment may be optimal

In clinical studies, the definitions of ‘short’ and ‘long’ antibiotic treatment vary [19, 29], and here we compare treatments that are either 3 (“short”) or 7 (“long”) days. The endpoint in clinical trials is typically a measurement of bacterial load or clinical symptoms at a given period after conclusion of therapy. We compare outcomes of 3- and 7-day treatment first on infection profiles (e.g. at the end of treatment), or using different metrics evaluated at 20 days post-infection.

By varying the duration of treatment and the time when treatment starts (defined by Ω), a variety of outcomes can be observed in the model (Figure 2); most importantly, both short and long treatments may result in treatment failure (defined as detectable bacteria at the end of treatment, e.g., Figure 2C&D). Similarly, both short and long treatments may be successful (e.g., Figure 2E&F). These results stem from the assumptions in the model [30] due to interplay between generation of the immune response (which aids in clearing the infection) and generation of resistance (which prevents clearance upon treatment). Specifically, longer treatment may help in faster clearance of bacteria when the treatment starts early (low Ω, Figure 2B) but leads to resistance selection and failure when treatment starts late, due to random generation of resistance strain by the time of treatment start (high Ω, Figure 2D &H). Shorter treatment resulted in bacterial clearance during treatment only when early and at higher dose (Figure 2E), but failing in the case of too low kill rates (Figure 2A). However, such antibiotic exposure was still able to limit bacterial growth post-treatment and constrain resistance selection, when applied later and moderately (Figure 2C). When higher antibiotic kill rates are applied later, short and long treatments result in similar dynamics of resistance selection during and post-treatment (Figure 2G&H).

**Figure 2:**
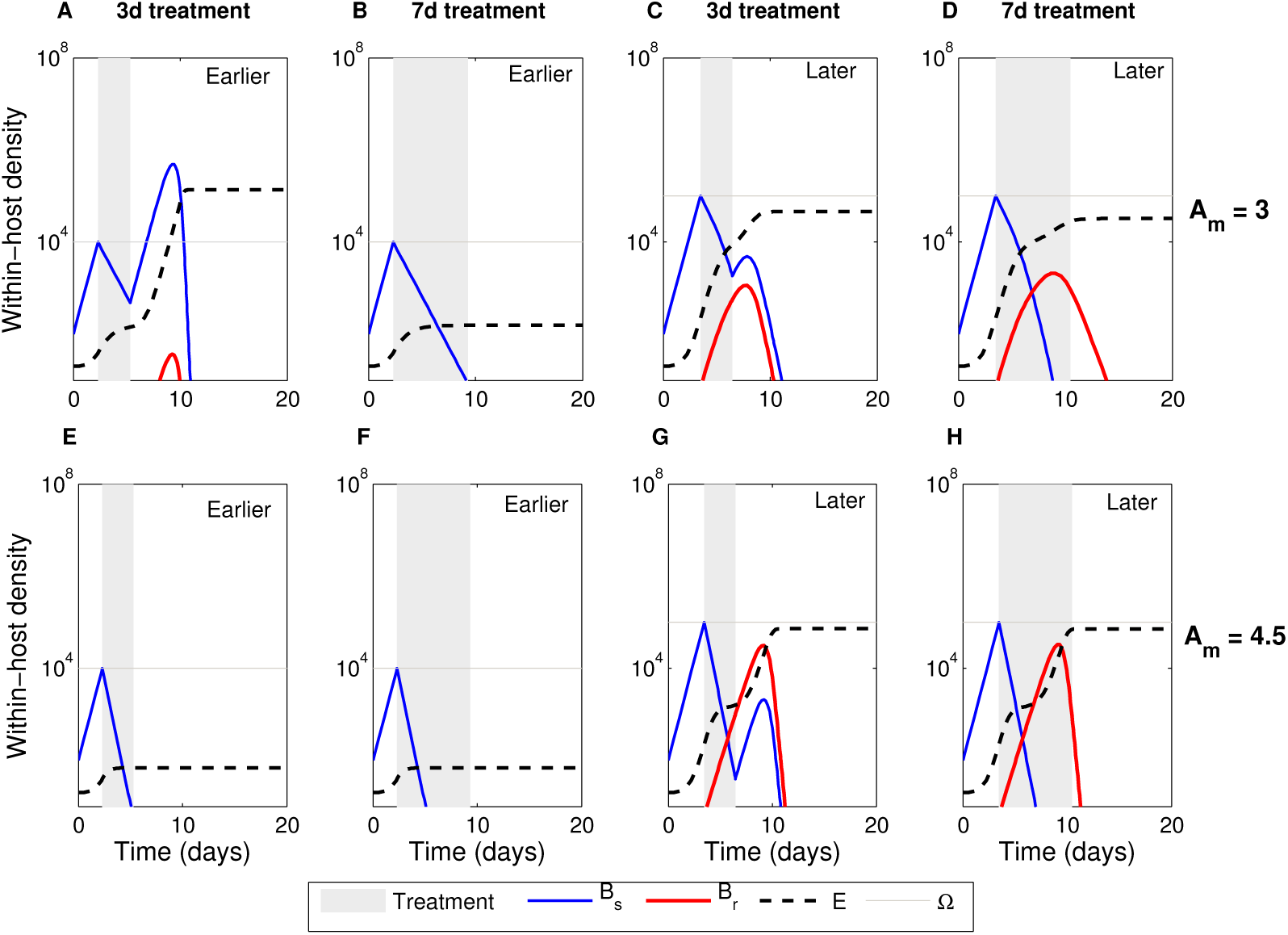
Both short and long treatments can result in treatment failure. Earlier onset: Ω = 10^4^, Late onset: Ω = 10^5^. We show dynamics with a short (3-day) and long (7-day) treatment for two antibiotic efficacies *A*_*m*_ = 3 **(A-D)** and *A*_*m*_ = 5 **(E-H)**, both representing supra-inhibitory levels. Parameters as specified in Table 1.

Another way of looking at this continuum of scenarios is by distinguishing treatment failures on one extreme where resistance is causative of the relapse (Figure 2D & H), and treatment failures unrelated to resistance per se (Figure 2A & C). In the first case, the relapse is due to slowed down immune response during treatment. In the second case, if resistance weren’t there, treatment would have resulted in the clearance of infection.

A more comprehensive analysis further demonstrates that optimal duration of treatment strongly depends on the chosen metric defining successful treatment, threshold of treatment start, and antibiotic kill rate (Figure 3). Two features are i) the non-monotic relation to antibiotic kill rate, and ii) the tendency to shorter non-inferior regimens, when higher antibiotic kill rates are applied. In particular, while 3-day treatment at moderate kill rate (*A*_*m*_ = 2/day) may be superior when started early when success of treatment is defined by the duration of infection (Figure 3A), it is inferior when success of treatment is measured by final bacterial burden (Figure 3D). Furthermore, even though 3-day treatment leads to a higher final immunity overall, the difference between the 3-day and 7-day treatment strongly depends on the antibiotic efficacy and timing of treatment, and it vanishes when treatment is started at higher pathogen loads (Figure 3I). Finally, while final immunity may be higher after a 3-day treatment (Figure 3H) at moderate kill rates, the corresponding values of cumulative immunopathology are lower than a 7-day treatment (Figure 3K), suggesting conflicting optima by instantaneous and cumulative indicators of success.

**Figure 3:**
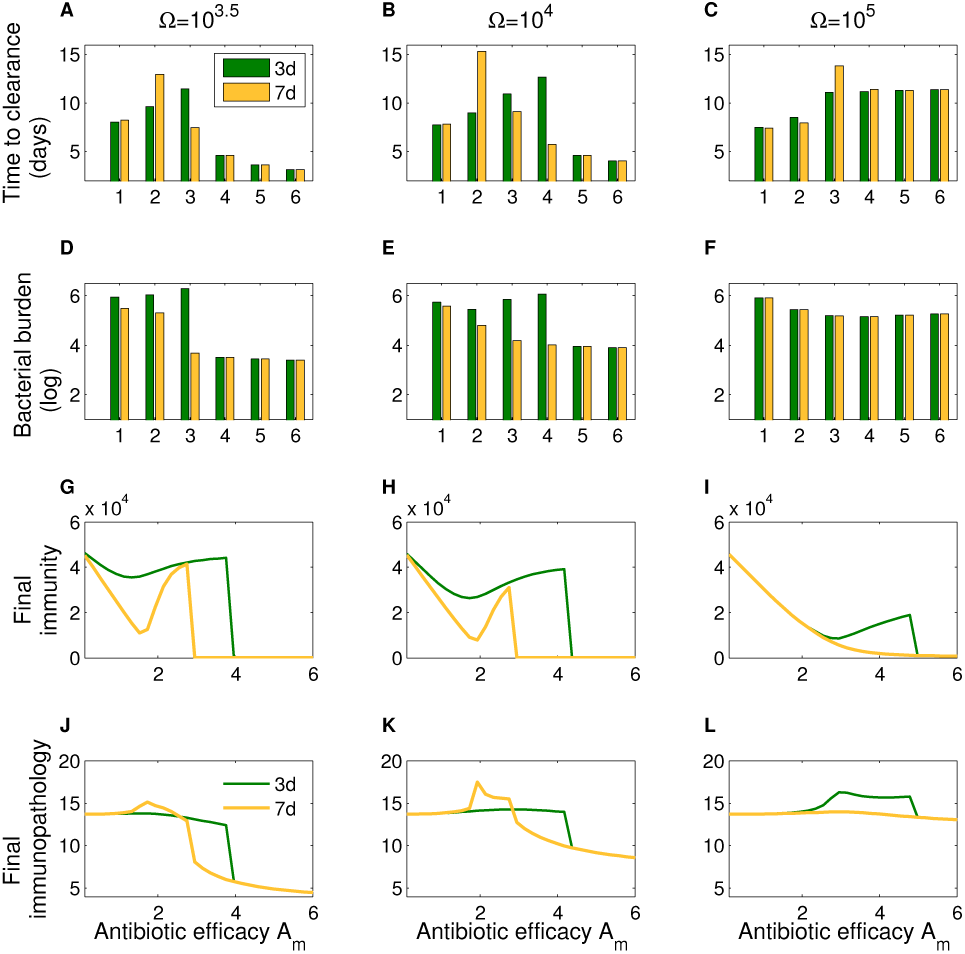
Benefits of short or long treatment depend on the metric of success.. We compare outcome of 3- and 7-day treatments for different metrics for successful treatment and antibiotic efficacy. Success of treatment is defined in terms of: **(A-C)** duration of infection, **(D-F)** bacterial burden, **(G-I)**, the final level of immunity, or **(J-L)** cumulative immunopathology level (Eqn. (4)). Infection was simulated for 20 days with default parameters (Table 1).

We further highlight the inability to select universally optimal duration of treatment by a series of simulations in which we varied duration of treatment and antibiotic efficacy continuously, for different timing of treatment, and measured deviation of several metrics of successful treatment from their baseline values (Supporting Information Fig. S1). We observe that the quantitative landscapes of different metrics are variable, thus confirming that each effectiveness indicator displays its own nonlinear dependence with duration, intensity, and timing of treatment. Furthermore, as the sensitivity of these metrics is typically higher for the antibiotic efficacy axis, the more beneficial treatments tend to include short and long treatments where crucially, the same effects can be achieved using minimally effective therapeutic durations. When treatment onset occurs later during the infection course, unsurprisingly, the marginal benefits of treatment decrease for most metrics, making reductions of at least 50% in target criteria relative to no treatment harder to reach. However such later timing appears to shift the optimality region to high-antibiotic efficacy and short duration combinations, especially for resistance selection.

### Treatment timing and the immunity-resistance trade-off

Typically the timing of treatment relative to the natural infection course is not known. Previous models assumed that treatment occurs throughout infection or when infection reaches a peak [23, 22], which imposed strong constrains on the model dynamics. We have shown that relaxing this assumption and starting the treatment at different bacterial densities results in dramatically different predictions on which specific treatment duration is optimal.

In this model, the key to understanding how treatment duration affects infection dynamics is how bacterial load impacts generation of the immune response. We assumed that immune response expansion is directly driven by the amount of bacteria and that immune response does not contract during the timescale of infection. This implies that the net growth rate of bacteria is dynamic: it declines progressively due to the activation of the immune response, until it reaches negative values for super-critical levels of immunity (Figure 4). In parallel, due to increasing bacterial levels, the probability of resistance emergence increases. Nonetheless, this increase in chances of emergence is counterbalanced by the top-down immune control, whereby the effective growth potential of any resistant sub-populations gets progressively reduced, limiting their ascent and eventual competitive release via treatment. In this context, any treatment that does not result in clearance of bacteria but significantly slows down immune response dynamics, may then lead to relapse. And the later such treatment is administered, the higher the probability that this relapse is constituted by a majority of resistant bacteria, freed by their competitors, and favored by weakened immunity.

**Figure 4:**
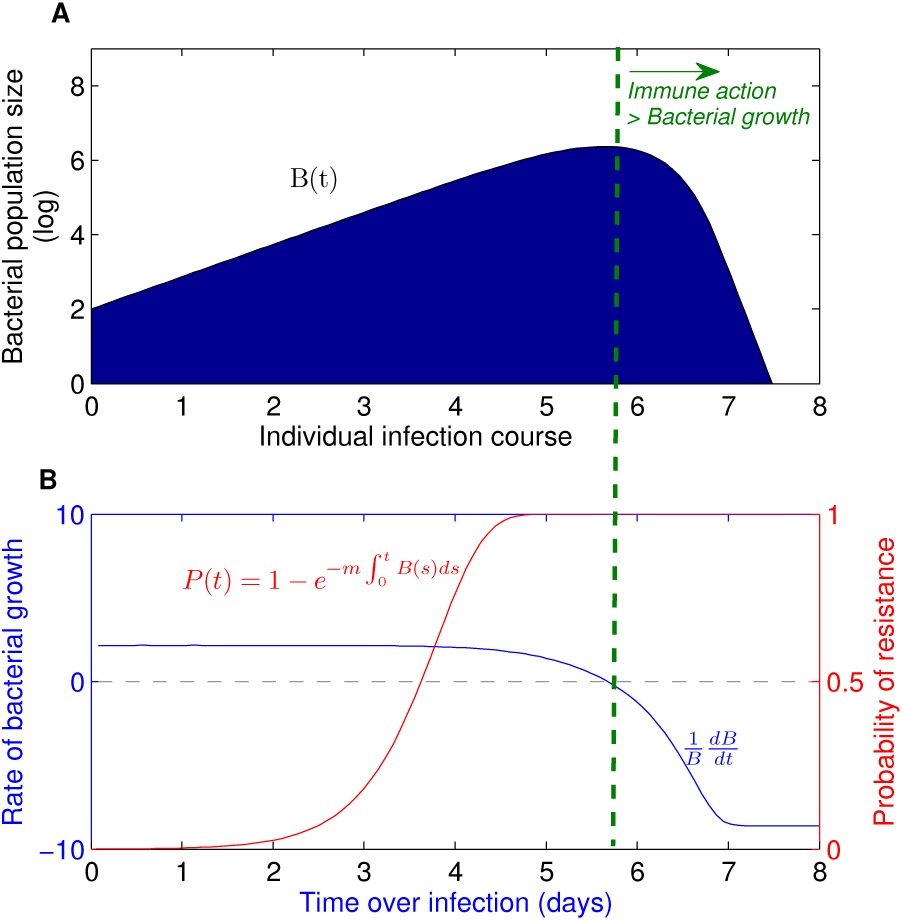
Immunity-resistance trade-off. **A**. Graphical illustration of typical acute infection dynamics. **B**. Probability of resistance emergence (*P*(*t*)) increases over infection, while the net rate of bacterial growth 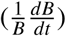 decreases as a result of immune activation. Impairment of immune buildup at later stages of infection could therefore select for more resistance.

Such trade-off between immunity and resistance is naturally modulated by treatment onset, fitness cost of resistance, and activation kinetics of immune response, which ultimately determine how fast immunity grows and antibiotic resistance ascends, and which force is strongest in a given infection. Although the quantitative details likely vary with parameter values, qualitatively, we find that this trade-off can lead to opposite trends between resistance selection and treatment duration, depending on treatment start (Figure 5). In particular, when treatment happens relatively early over the infection course, longer treatments tend to reduce resistance selection. In contrast, when treatment happens relatively later, the situation gets reversed: longer treatments tend to increase resistance selection. This negative association trend for early treatment becomes even more pronounced and significant when stronger antibiotic kill rates are used (Supporting Information Fig. S2), while the positive association trend between duration and resistance selection in later treatment is reinforced by lower immune competence of the host (Supporting Information Fig. S3). It should be noted, however, that absolute resistance levels increase when treatment starts late (e.g., Figure 5A vs. Figure 5C), simply because of a higher chance of resistance being present later in infection (Figure 4). A more comprehensive analysis of Spearman correlation coefficients (Supporting Information Fig. S4) confirms the treatment-timing effect on the monotonic relationship between resistance levels and length of antibiotic exposure.

**Figure 5:**
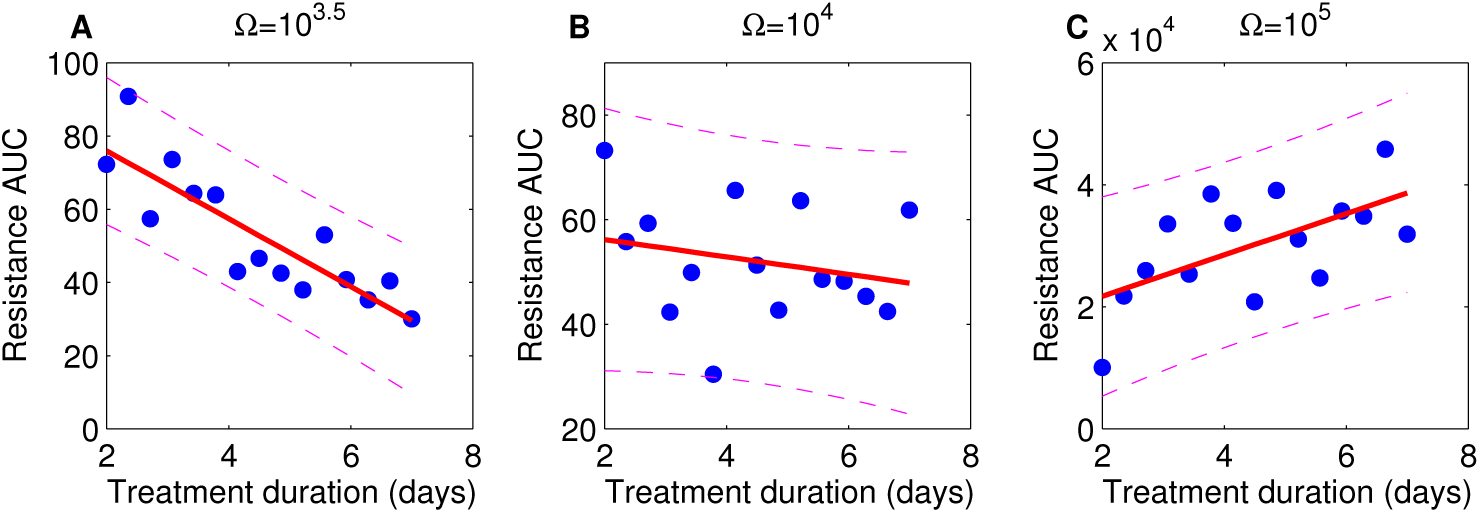
Treatment timing changes the trend between resistance selection and treatment duration. We simulated 30 treatments with random values of kill rates *A*_*m*_ ∈ [0.1, 6] for each treatment duration *τ* ∈ [2, 7] days, and computed the overall resistance burden (AUC_*R*_) for early (**A**, Ω = 10^3.5^), intermediate (**B**, Ω = 10^4^), or late (**C**, Ω = 10^5^) treatment start times. Markers indicate mean values over all kill rates for each treatment duration (parameters as in Table 1), and the red line a linear fit *y* = *a* + *bx* in Matlab, together with 95% CI (dashed lines) for illustration. Spearman correlation coefficients: A. Negative trend (*ρ* = − 0.9, *p* < 0.01) B. Negative trend (*ρ* = − 0.24, *p* = 0.4) C. Positive trend (*ρ* = 0.52, *p* < 0.05). A more general analysis of correlative trends for the monotonic relationship (multiple Spearman correlation coefficients) and their significance, across many realizations such as this one, is shown in detail in Supporting Information Fig. S4.

Considering that typically infected hosts seek treatments at later stages of their infection, the notion that higher antibiotic exposure via longer treatments leads to higher levels of resistance, would favor shorter antibiotic courses in most cases. While the symptomatic causes of treatment remain often unclear, our results suggest that in less tolerant hosts who, instead, seek treatment at lower pathogen thresholds, longer antibiotic courses would be better to constrain resistance risk.

## Discussion

Antibiotics are key to modern medicine and preserving their effectiveness is a global priority. Avoiding antibiotic overuse remains an essential step in addressing the antibiotic resistance challenge [31, 32, 9]. Epidemiological studies have linked higher antibiotic exposure to higher levels of antibiotic resistance [33, 34], sometimes with contrasting results [35]. It is likely that a better understanding of how individual infection processes and treatment parameters affect resistance dynamics will help devising efficient treatment strategies that use minimal drug amounts. In this study, we focused on a comparison of treatment outcome between short and long antibiotic treatment, taking into account several treatment target criteria, and highlighting a previously neglected parameter of treatment start time (but see [27, 36, 37]). We found several parameter regimes and target criteria by which 3-day treatment is non-inferior and even superior to 7-day treatment, favoring a reduced use of antibiotics to achieve similar clinical outcomes. However, we also observe that optimal treatment duration varied widely, depending on the time point of treatment, antibiotic efficacy, host immunity characteristics, and the treatment criterion to optimize.

This difficulty in drawing general principles from a multi-factorial problem is not new. At the epidemiological level, it is also being recognized that ranking antibiotic treatment protocols is highly dependent on methodological factors, e.g. the criterion of choice for comparison [38]. In studies of the evolution of virulence, early work has argued that a single factor such as the route of transmission determines virulence of pathogens [39, 40], while later studies showed that a single factor is unlikely to determine optimal virulence of very diverse pathogens [41]. Hard-to-reach conclusions on optimality of antibiotic treatment duration might tempt one to logically constrain mathematical models so that a robust conclusion can be reached. For example, by assuming that immune system-mediated killing of bacteria is always high, treatment does not have to be long to be efficient since strong immunity can easily finish the job of eliminating bacteria if antibiotic does not [25]. However, what makes immune response “strong”, how quickly it becomes “strong”, how long it is “strong” for, and how the strength varies between individuals remains unknown, challenging the applicability of such a generic argument for the actual bacterial infections of humans. Better understanding of optimal duration of treatment necessarily requires solid empirical data, which can constrain theoretical models and may allow to discriminate between alternatives [30].

There are a number of potential limitations of our modeling analysis. We did not study the role of transmitted pre-existing resistance; starting the infection with already pre-existing resistant variants is likely to disfavor longer treatments [14]. We ignored pharmacokinetics and pharmacodynamics of the antibiotics which may also on their own add uncertainty in defining optimal treatment duration. We considered a single mutational step to complete resistance. This is likely a worst-case scenario. Resistance may be acquired in multiple gradual mutational steps [4, 42] where transient phenotypes in fitness cost and antibiotic susceptibility may interact differently with treatment, slowing down evolution.

Our mathematical model is broadly deterministic with a hybrid component in emergence of resistant bacteria, similar to the deterministic threshold approximation used previously (e.g., [43]). Changing the threshold for probability of emergence to random values confirmed that the behaviors described in the paper capture up to 80% dynamics under stochastic arrival times for the resistant strain (Supporting Information Figs. S5–S7). Furthermore, this variation did not influence the overall conclusion that selection of optimal treatment duration strongly depends on infection details (Supporting Information Fig. S8).

Perhaps the strongest limitation of the model is its lack of specificity and exact parameter estimates for bacterial infections of humans from which to derive final quantitative conclusions. This limitation is common for most (if not all) of the previously published mathematical modeling-based studies. Our approach was to consider perhaps the simplest mathematical model of a bacterial infection which includes resistance to antibiotics and control by host immunity; even such a simple model already generates a variety of complex behaviors precluding reaching a solid conclusion on the uniformly optimal antibiotic treatment duration. Conceivably, use of more complex mathematical models with more poorly constrained parameters is likely to display even richer behaviors.

Model predictions are logical extensions of the assumptions, and therefore, if the list of assumptions is unbounded, any conclusions can be reached [30]. Experimental studies by measuring dynamics of infections and infection-controlling immune responses in humans will allow to constrain the model assumptions — to define which assumptions are unrealistic — and thus, to build more accurate mathematical models. Such constrained models are more likely to predict when a treatment will succeed and when it will fail. Testing such model predictions in clinical trials will be important, especially when anticipated outcomes do not match the observations [44].

What are the unknown details of bacterial infections of humans? There are many. For example, the kinetics of most acute bacterial infections in humans have not been accurately measured (but see [45, 46]). One critical parameter in our analysis was the time when treatment starts (which in the model was strictly determined by the bacterial density). Physiological factors driving patient symptoms and treatment onset remain unclear, and are likely to widely vary between individuals, as evidenced, for example, by differences in microbiologic confirmation at baseline across patients with the same symptoms [29].

Furthermore, how the dynamics of immune response, both innate and adaptive, depends on the presence of the infection in humans is not understood and may well depend on the type of infection [47] and different aspects of host susceptibility [48]. In particular, yellow fever virus-specific CD8 T cell response continues to expand long after the infection disappears from the blood [49, 50]. How immunity clears the infection is also unknown, and, for example, if killing of drug-sensitive and drug-resistant bacteria by the immune response occurs at the same rate – an assumption made by most previous studies (e.g., [27, 25]). Unequal distribution of antibiotics in different tissues or tissue compartments may select for resistance at different rates, effectively resulting in different rate of bacteria elimination by immunity, especially if antimicrobial susceptibility and susceptibility to host immunity correlate.

Better understanding of mechanisms regulating abundance of bacteria and bacteria-specific immune responses in tissues in human infections is likely to constrain mathematical models, and will help to make more robust predictions about eventual intervention strategies.

## Materials and methods

### Simplest mathematical model of a bacterial infection

We extend a previous model of an acute infection controlled by the immune response [28] by allowing generation of antibiotic-resistant strain and by including antibiotic treatment. Infection is initiated by the drug-sensitive bacteria (*B*_*s*_) which grow exponentially at a rate *r* until reaching a carrying capacity *K* (i.e., the bacterial dynamics follows a logistic growth as it has been observed in humans, e.g., [46]). Drug-resistant bacteria (*B*_*r*_) are generated via mutation with rate *m*; thus, the total density of bacteria in the host is *B* = *B*_*s*_ + *B*_*r*_. For resistance emergence, we adopt a hybrid approach, tracking the probability of no-emergence by time *t*, 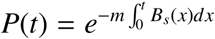, and simulating emergence of the resistant sub-population when the probability hits a deterministic threshold *P* = 0.5, similar to [43]. *B*_*r*_ at that emergence time point, *t*_*em*_, are initialized at the value *B*_*r*_(*t*_*em*_) = *B*_*em*_, and subsequently let to follow deterministic growth. Generation and extinction are handled through the ODE Events option in Matlab ode solver ode45. We assume an extinction threshold *B*_*ext*_ when either bacterial compartment falls below this level. We assume that resistance bears a cost *γ* which reduces the growth rate of the drug-resistant variant to *r*(1 − *γ*). Antibiotic treatment starts when the total bacterial density exceeds value Ω, lasts for *τ* days, and is described by the step-function η. Antibiotics increase the death rate of drug-sensitive bacteria *B*_*s*_ by the rate *A*_*m*_ (antibiotic kill rate), while the *B*_*r*_ strain is fully drug-resistant.

For bacterial infections of humans it is generally poorly understood which types of immunity – innate or adaptive – are most important in control of the specific infection. Here we do not make an explicit distinction between innate and adaptive immunity, and rather implement immune response kinetics generically via two mathematical features: antigen-dependent stimulation and negative feedback on infection. The major assumption we make is that immune response gets triggered after total bacteria reach some density, defined by the half-saturation constant *k*. When bacterial density is high, immune response magnitude increases exponentially at maximal rate σ until infection is cleared. This immune response eliminates sensitive and resistant bacteria at equal rate *d*. The initial level is given by *E*(0) = *E*_0_. With these assumptions, the model is given by the following ordinary differential equations:

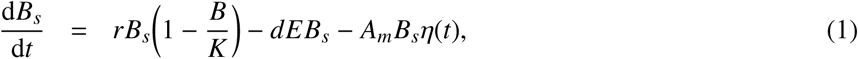

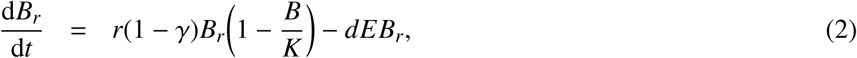

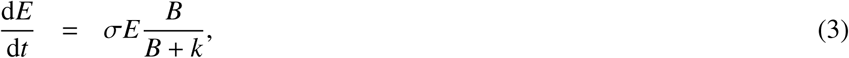

where *B* = *B*_*s*_ + *B*_*r*_ and *η*(*t*) is varied between 0 and 1 to reflect treatment interval from onset through its duration.

### Treatment outcome metrics

Success of the antibiotic treatment can be defined in many different ways. In our analysis we consider the following metrics: i) duration of infection *T* (the period from *t* = 0 to *t* = *T* when all populations of bacteria reach an extinction threshold *B*_*ext*_), ii) total bacterial burden 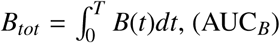, iii) total resistance burden 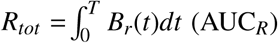, iv) final immunity *E*_*T*_, and v) cumulative immunopathology 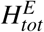, defined as the cumulative deviation from initial immunity:

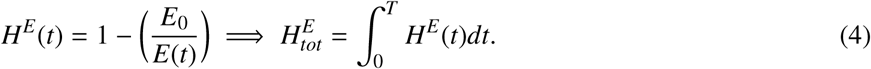

### Model parameters

The formulated mathematical model is likely to apply to extra- and intracellular bacterial infections and is also similar to previously proposed models for viral infections [51]. Quantitative details of bacterial infections of humans are nearly absent, and therefore, we chose model parameters to constrain the overall bacterial dynamics in the absence of treatment. Specifically, bacteria are likely to reduce their replication rate within a few days since infection [46] which in our model required rapidly activating and expanding immune response. To our knowledge, the kinetics of immune response to bacteria in humans has not been accurately measured, but virus-specific T cell responses tend to expand relatively slowly [49, 52, 50]. Antibiotic kill rates have been accurately measured for several drugs *in vitro* [53] but not in humans, and thus were varied within expected range.

## Acknowledgements

This research was supported by a NOS Alive-Instituto Gulbenkian de Ciência fellowship awarded to F.F.S.P. in 2017. The work was also in part assisted by NIH grant (R01 GM118553) to V.V.G and a FLAD/NSF grant (274/2016) to E.G. The National Institute for Mathematical and Biological Synthesis (NIMBioS) provided support through a short-term visitor grant to E.G. at the University of Tennessee, Knoxville.

## Author contributions

E.G. and V.V.G conceived and designed the study. F.P carried out initial numerical simulations and contributed to the interpretation of results. V.V.G provided critical feedback during all phases of the work. E.G. and V.V.G wrote the manuscript. E.G. supervised all technical and analytical details of the study. All authors have given final approval of the version submitted.

## Supplemental Information

**Supplemental Figure S1:**
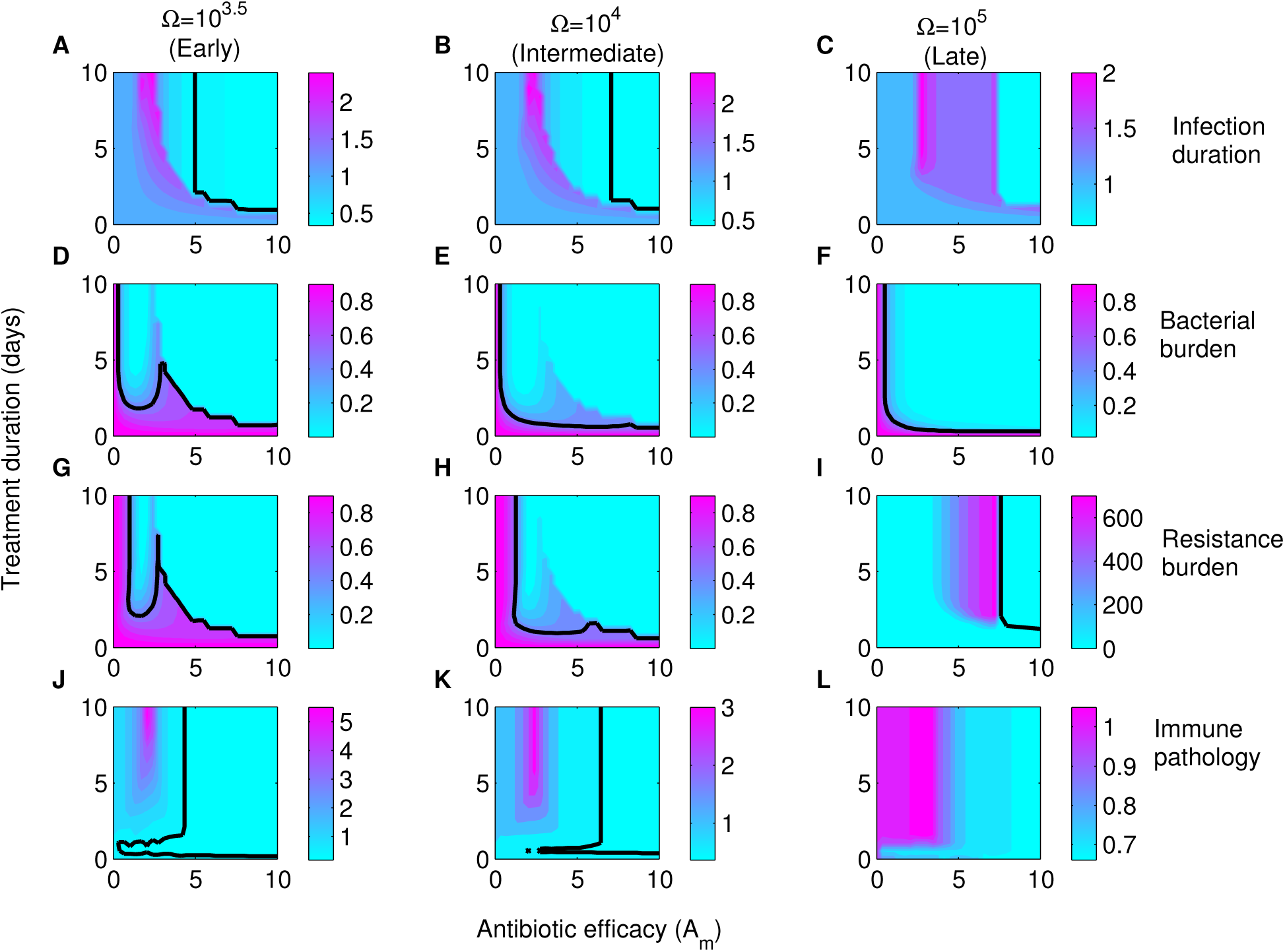
Intensity-duration treatment landscapes. The contour plots show quantitative treatment effects (ratios relative to no-treatment) on several infection metrics, varying antibiotic strength (*A*_*m*_) and treatment duration simultaneously. We examine three treatment timings to illustrate contrasting qualitative patterns: earlier treatment, when *B*(*t*) = Ω = 10^3.5^ (first column), intermediate timing *B*(*t*) = Ω = 10^4^ (second column) and later treatment when *B*(*t*) = Ω = 10^5^ (third column). **(A-C)** Infection duration. **(D-F)** Bacterial burden. **(G-I)** Resistance selection. **(J-L)** Immuno-pathology. The black solid line depicts the 0.5 contour line relative to no-treatment, when applicable.

**Supplemental Figure S2:**
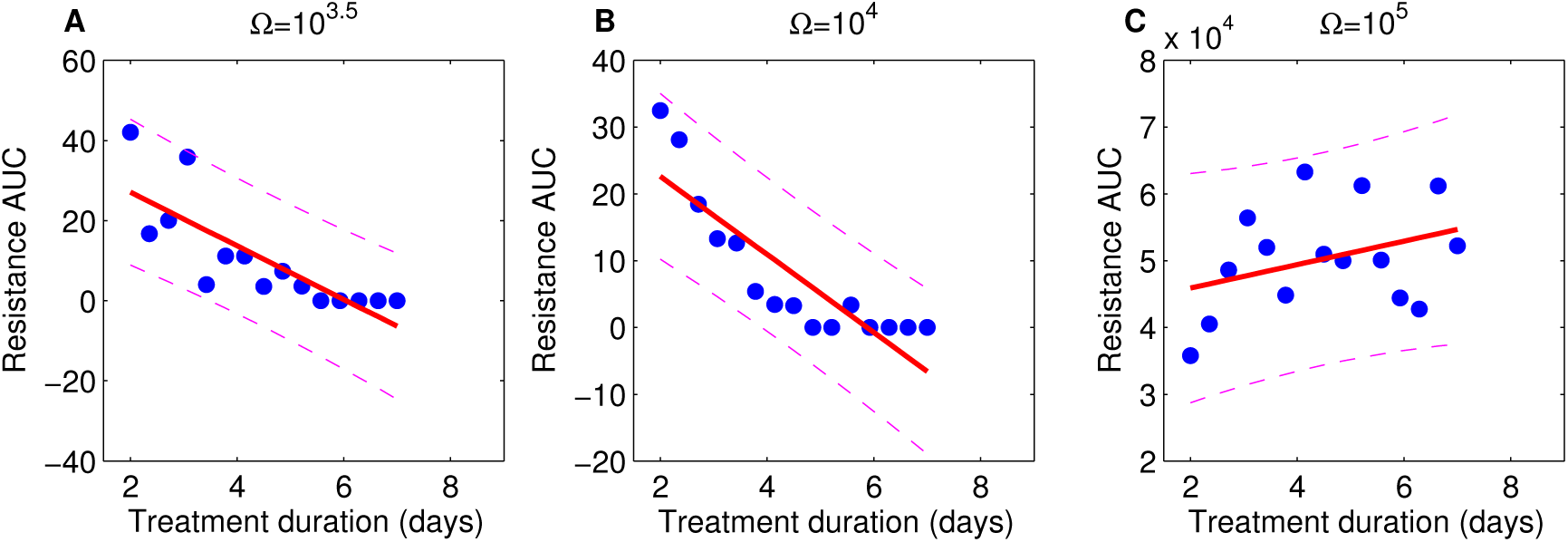
Resistance vs. treatment duration, over bactericidal doses. Replicate of Fig. 5. (each scatter point represents the mean *R*_*tot*_ of 30 simulations with random *A*_*m*_ ∈ [3, 6] for each duration *τ* ∈ [2, 7]). All parameters as in Table 1. The pattern of treatment onset-dependent trends in Figure 5 changes with more aggressive kill rates. Results of the linear fit *y* = *a* + *bx*, for each treatment timing, are plotted for illustration, while the Spearman correlation coefficients are: A. (*ρ* = −0.93, *p* < 0.001) B. (*ρ* = −0.93, *p* < 0.001) C. (*ρ* = 0.33, *p* = 0.22)

**Supplemental Figure S3:**
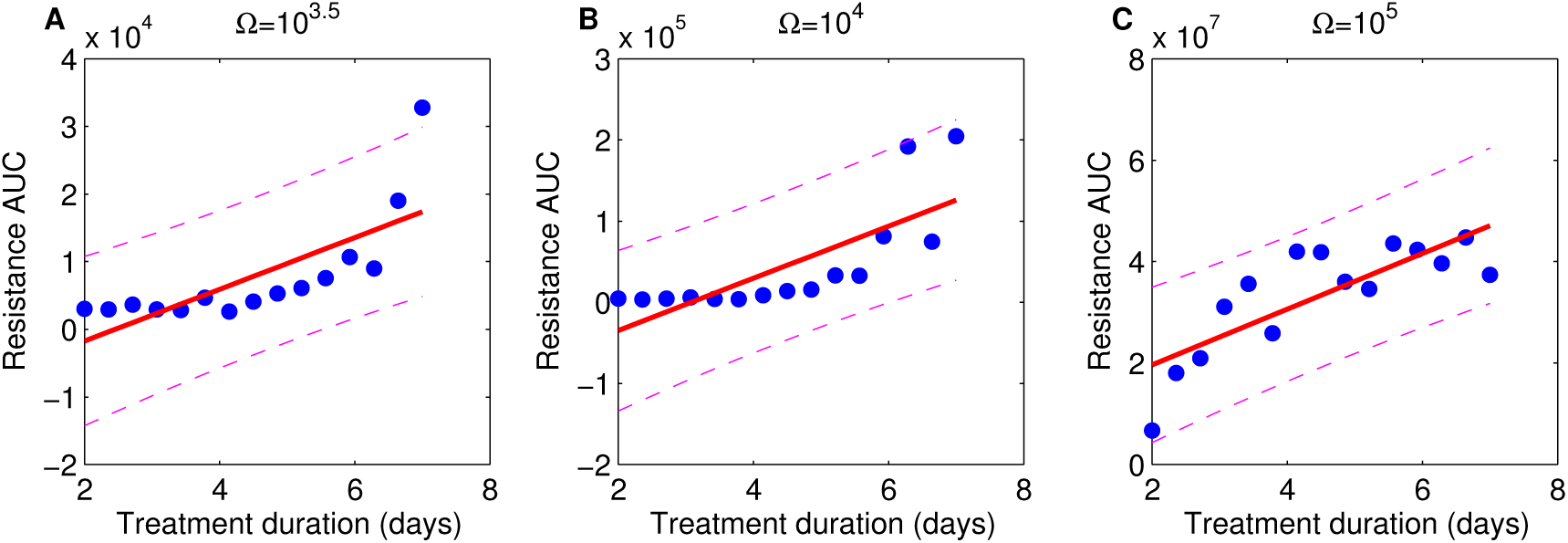
Resistance vs. treatment duration, in immune-compromised hosts (σ = 0.5). Replicate of Fig. 5. (each scatter point represents the mean *R*_*tot*_ of 30 simulations with random *A*_*m*_ ∈ [0.1, 6] for each duration *τ* ∈ [2, 7]). All parameters as in Table 1. In hosts with lower activation rate of the immune response as a function of pathogen density, longer treatment duration leads to increased resistance over more treatment onset scenarios. Results of the linear fit *y* = *a* + *bx* are plotted for illustration. Spearman correlation coefficients for trends are: A. *ρ* = 0.35, *p* = 0.19 B. *ρ* = 0.22, *p* = 0.41 C. *ρ* = 0.77, *p* < 0.01. The positive trend becomes significant in the late timing scenario.

**Supplemental Figure S4:**
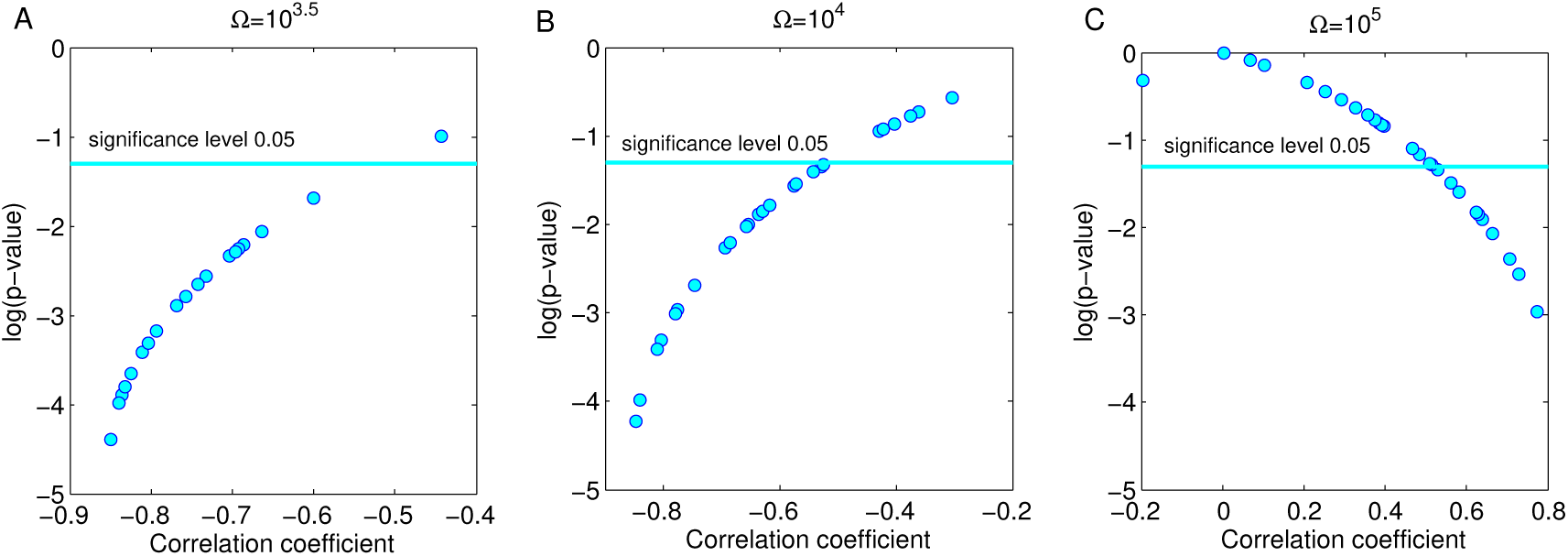
Correlation coefficient for the association between resistance selection and treatment duration. 30 random realizations, each replicating the data in Fig. 5, are shown. Here, each scatter point represents the Spearman correlation coefficient and corresponding p-value, between mean *R*_*tot*_ (over 30 simulations with random *A*_*m*_ ∈ [0.1, 6]) and treatment duration *τ* ∈ [2, 7]. All parameters as in Table 1. A) Early treatment (Ω = 10^3.5^ results in a negative and significant association between resistance selection and treatment duration. B) Intermediate treatment onset (Ω = 10^4^) results in negative, but less strong associations between *R*_*tot*_ and treatment duration. C) Later treatment (Ω = 10^5^) results in positive and generally significant associations between resistance and treatment duration.

**Supplemental Figure S5:**
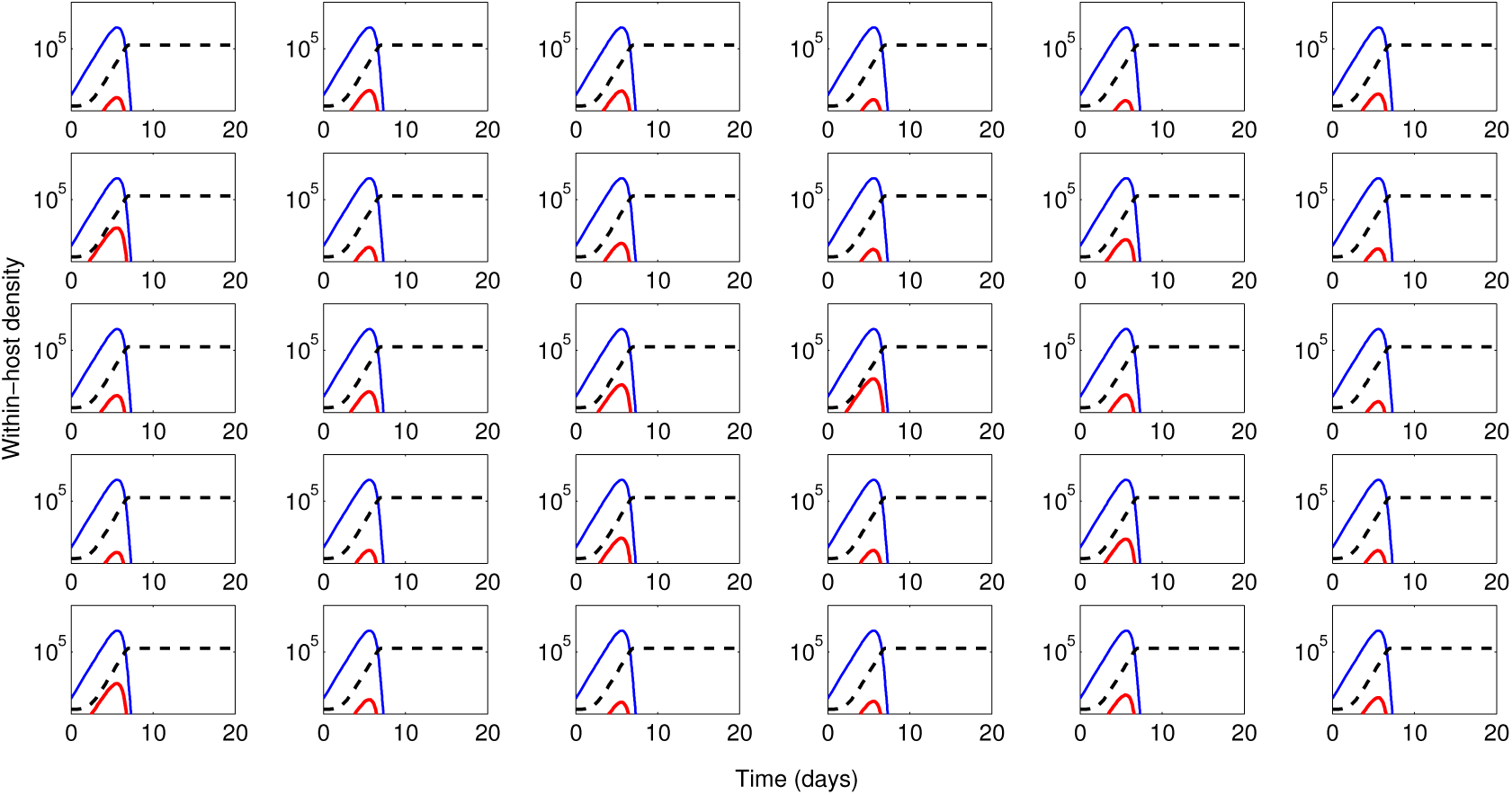
Model dynamics for random arrival times of the resistant mutant (30 realizations). All parameters as in Table 1. The threshold for probability of emergence is set to a random uniform variable between 0 and 1, instead of the fixed number 0.5, assumed in the paper. Replicates of Fig.1A.

**Supplemental Figure S6:**
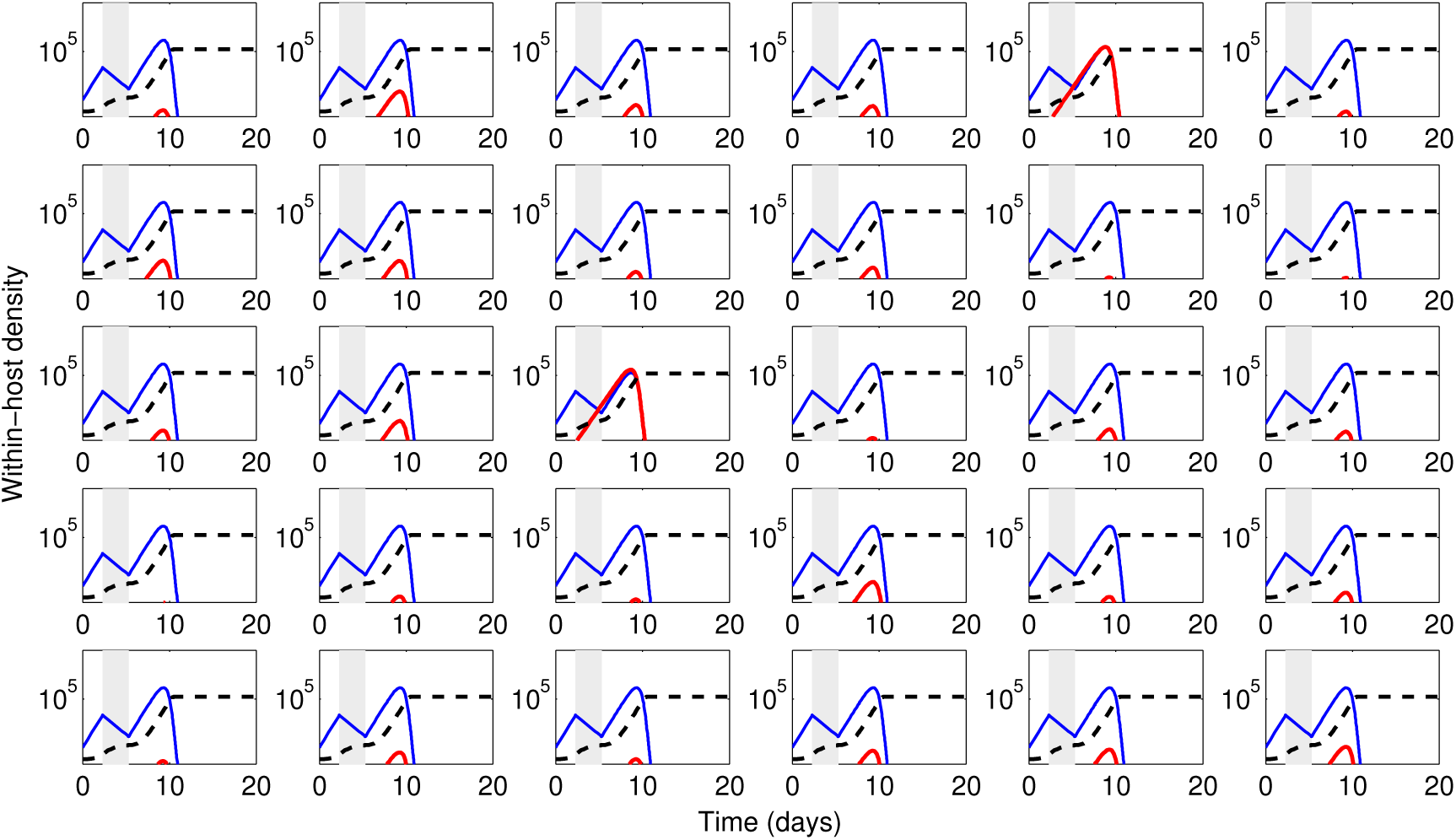
Model dynamics with treatment (*A*_*m*_ = 3, *τ* = 3, Ω = 10^4^) for random arrival times of the resistant mutant (30 realizations). All parameters as in Table 1. The threshold for probability of emergence is set to a random uniform variable between 0 and 1, instead of the fixed number 0.5, assumed in the paper. Replicates of Fig. 2A. Only in 2 out of 30 cases (< 10%) the short treatment resulted in qualitatively different dynamics (resistance selection) than the one captured by the average deterministic threshold approximation. This due to the rare event of extremely early stochastic emergence of *B*_*r*_.

**Supplemental Figure S7:**
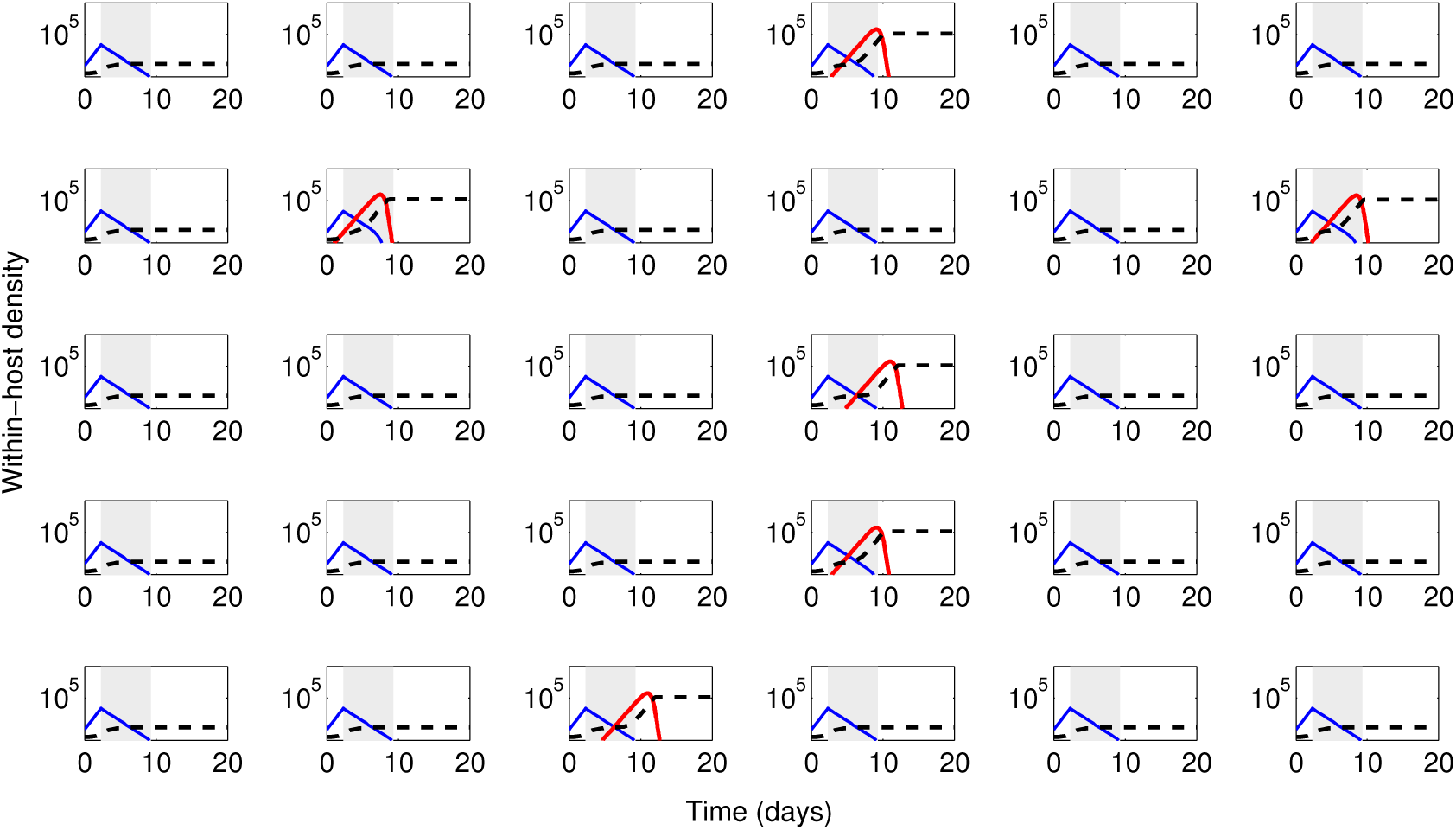
Model dynamics with treatment (*A*_*m*_ = 3, *τ* = 7, Ω = 10^4^) for random arrival times of the resistant mutant (30 realizations). All parameters as in Table 1. The threshold for probability of emergence is set to a random uniform variable between 0 and 1, instead of the fixed number 0.5, assumed in the paper. Replicates of Fig. 2B. Only in 6 out of 30 cases (=20%) the long treatment resulted in qualitatively different dynamics (resistance selection and longer infection) than the one captured by the average deterministic threshold approximation. This due to the rare event of extremely early stochastic emergence of *B*_*r*_.

**Supplemental Figure S8:**
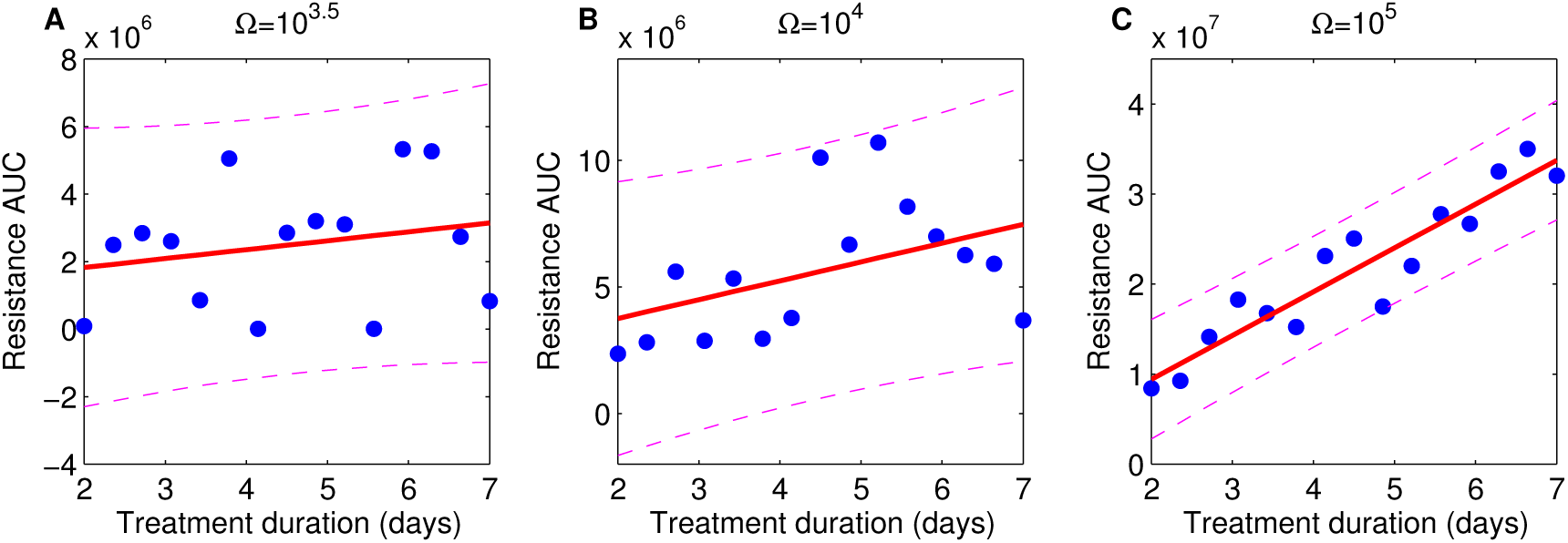
Resistance vs. treatment duration, in immune-compromised hosts (σ = 0.5) with stochastic arrival times of resistance. Replicate of Fig. 5. (each scatter point represents the mean *R*_*tot*_ of 30 simulations with random *A*_*m*_ ∈ [0.1, 6]) for each duration *τ* ∈ [2, 7]. All parameters as in Table 1, but here instead of a fixed threshold for stochastic emergence, a random threshold is used in each model simulation. Linear fits are shown for illustration of trends. Spearman correlation coefficients are: A. *ρ* = 0.26, *p* = 0.4 B. *ρ* = 0.57, *p* < 0.05, C. *ρ* = 0.92, *p* < 0.01. Compared to Figure S3, the same qualitative trends are preserved under more stochasticity in resistance arrival times.

